# Biological and cultural drivers of oral microbiota in Medieval and Post-Medieval London, UK

**DOI:** 10.1101/343889

**Authors:** A. G. Farrer, J. Bekvalac, R. Redfern, N. Gully, K. Dobney, A. Cooper, L. S. Weyrich

**Affiliations:** Australian Centre for Ancient DNA, School of Biological Sciences, University of Adelaide, Adelaide, South Australia, Australia; Centre for Human Bioarchaeology, Museum of London, London, United Kingdom; School of Dentistry, Faculty of Health Sciences, University of Adelaide, Adelaide, South Australia, Australia; Department of Archaeology, Classics and Egyptology, School of Histories, Languages and Cultures, University of Liverpool, Liverpool, United Kingdom; Department of Archaeology, University of Aberdeen, Aberdeen, United Kingdom; Department of Archaeology, Simon Fraser University, Burnaby, Canada

**Keywords:** ancient DNA, dental calculus, microbiota, microbiome, Britain, London

## Abstract

The trillions of microorganisms that live in association with the human body (microbiota) are critical for human health and disease, but there is a limited understanding of how cultural and environmental factors shaped our microbiota diversity through time. However, biomolecular remnants of the human oral microbiota - recovered from the calcified dental plaque (calculus) of our long-dead ancestors - are providing a new means of exploring this key relationship of our evolutionary history. Here, we correlate extensive experimental, archaeological, and biological metadata with 128 ancient dental calculus specimens from Medieval and Post-Medieval London, UK (1066 – 1853 CE). We identify a significant association between microbiota and oral geography (*i.e.* tooth type and tooth surface), which has confounded ancient microbiota studies to date. By controlling for oral geography, however, we identify the first associations between ancient microbiota and cultural and environmental signatures. We find significant links between ancient British microbiota structure and health, including skeletal markers of stress that may reflect low socioeconomic status. Furthermore, this study provides baseline data to explore factors that drive microbiota differentiation within and between ancient populations and highlights the potential of ancient microbiota to infer detailed health and sociocultural information about the past.

## Introduction

It is now widely recognized that microbial communities both on and in the human body (microbiota) fulfill key functional roles that include accessing nutrients otherwise inaccessible from food resources, removing dead epithelial cells, restoring tooth enamel, and interacting with the immune system (1–5). Conversely, many diseases have also been linked to alterations in microbiota composition and/or function, including oral disease, arthritis, respiratory disorders, cancer, obesity, mental disorders, and many more (6–10). While several studies have examined how modern commensal bacterial communities respond to changes in lifestyle, diet, and environment (11–13), we know almost nothing about their evolutionary history or how that history shaped our own bio-cultural identity.

Bioarchaeological remains contain key data from natural experiments run in the past that can reveal the history of the bacterial communities found in modern populations, including the effects of different living conditions, diet, and disease (14–17). The analysis of such data is now routinely possible using ancient DNA (aDNA) recovered from calcified dental plaque (calculus), which is widespread in the archaeological record (18). Dental calculus is a calcified microbial biofilm on the surface of the teeth (19). This matrix preserves human microbiota in real-time during the life of the individual, forming the only reliable record of its kind in the archaeological record (15,18,20–23). Bacterial DNA recovered from ancient dental calculus has already revealed major changes in human oral microbiota across the major human biocultural transitions of the Holocene/Anthropocene (~10,000 ybp and ~200 ybp, respectively) (18), as well as information on real-time pathogen evolution (22,23). Eukaryotic DNA recovered from ancient dental calculus has also revealed differences in diet between past populations (22,23), However, no detailed, longterm studies of ancient microbiota within a single human population through time exists and, as such, many cultural and environmental factors that shaped our ancient microbiota remain unknown.

In addition, several fundamental technical issues remain unresolved in ancient dental calculus research. For example, the influence of oral geography has not been assessed (*i.e.* if differences between the specific sampling location in the mouth significantly impact results). Microbial composition is known to vary between tooth surfaces in modern individuals (24), but this is yet to be explored in detail in studies of ancient oral microbiota. In addition, recent studies of ancient dental calculus have been limited to very small sample sizes across large geographic ranges, likely limiting the ability to detect specific factors that shape ancient microbiota. Lastly, contamination control and detection is a critical issue, especially in samples with low levels of endogenous DNA (such as ancient dental calculus). In such samples, background or contaminant DNA from the laboratory can easily be high enough to drive signals (25). Filtering methods can be used to conservatively remove contaminant species (23); however, there is not currently an accepted systematic approach to wholly assess endogenous oral signal within dental calculus. Such unresolved issues might easily provide false positive results or confound real historical patterns, and these issues will become more important as studies seek to increase resolution.

Here, we examine the largest number of ancient dental calculus samples studied to date – from 128 Medieval and Post-Medieval Londoners [1066 – 1853]. We focused on a single city in order to remove geographic variation and to explore the localised cultural and environmental factors and how they shaped the British oral microbiota. The dental calculus samples utilized for this study are part of the Museum of London’s (MoL) human osteology collections, which include extensive, additional and detailed information about dating, paleopathology, and cultural context (*e.g.* age, sex, diet, location, oral health, and systemic disease). We go on to combine this extensive archaeological and biological metadata with detailed experimental information in order to explore whether tooth type and location in the oral cavity significantly bias the conclusions drawn.

## Methods

### Ethics

Research was conducted after approval from the Human Research Ethics Committee at the University of Adelaide (H-2012-108) following agreement to sample from the MoL Research Committee.

### Archaeological Context and Site Information

160 samples were collected from nine archaeological sites in a 10-square mile section of London (16km^2^), which formed a continuous temporal sequence from 1066 – 1853 CE (Figure 1; Table S1). The datasets included burial sites and cemeteries of monastic (Bermondsey Abbey, Spital Square, Merton Priory, St. Benet Sharehog, St. Mary Graces, and St. Brides) and laymen (Guildhall Yard), low class (Cross Bones), and upper class (Chelsea Old Church). All individuals were over 18 years old and had extensive metadata collated in the MoL’s Wellcome Osteological Research Database (WORD), including sex and age estimates, blood disorders, dental and vertebral anomalies and pathologies, and joint disease (Table S1).

**Figure 1:**
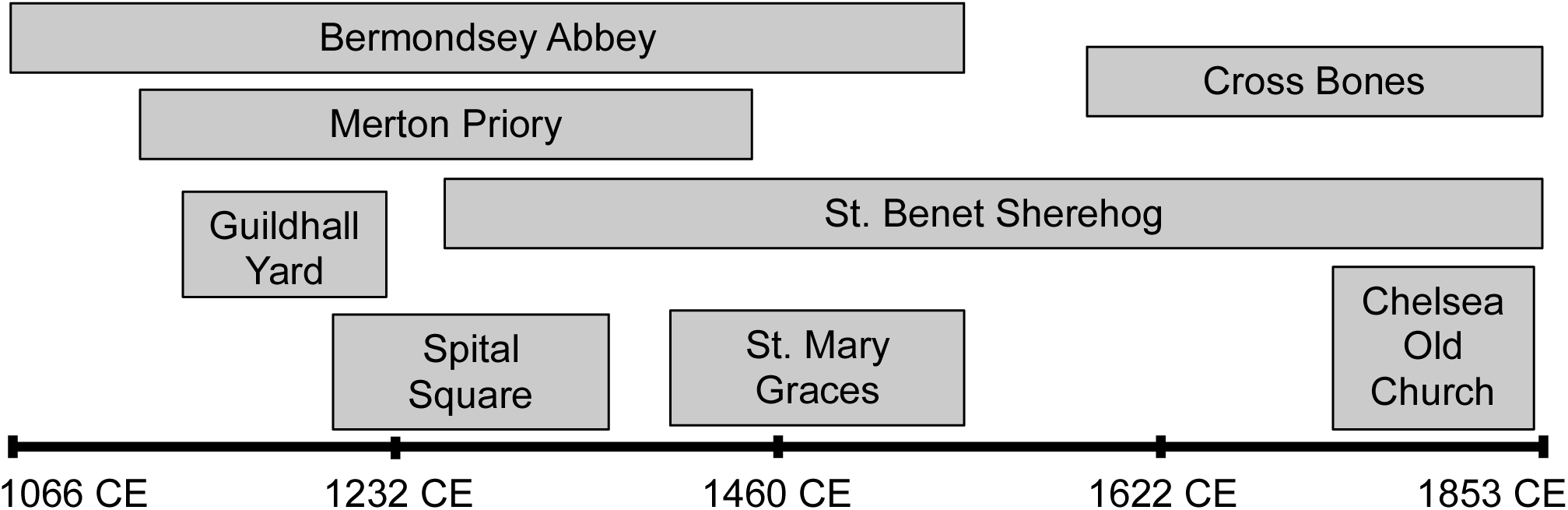
The eight archaeological sites included in this study and the length of time each site was actively used for burials. Samples throughout each of these periods are included in this study.

### Sample collection

All sampling was completed at the Museum of London using sterile procedures as previously published (14). Briefly, a facemask and gloves were worn to limit contamination from the researcher. The gloves were changed between each sample to limit cross-contamination. A sterile dental pick was used to remove the dental calculus deposit from a single surface of one tooth per skeleton. Pressure was applied in parallel to the tooth surface to ensure the enamel was not damaged. Calculus fragments were collected in sterile aluminum foil and placed into sterile plastic bags for transport to the aDNA facility at the Australian Centre for Ancient DNA, University of Adelaide, Australia.

### Sample decontamination, DNA extraction, shotgun library preparation, and DNA sequencing

All laboratory work was conducted in the specialized stand-alone aDNA facility at the Australian Centre for Ancient DNA at the University of Adelaide. Samples were processed in a random order to avoid bias. All calculus samples were decontaminated with bleach and UV exposure, as described elsewhere (18). To recover preserved DNA, dry samples were powdered in a sterile tube, immediately following decontamination. A modified silica-based DNA extraction was used on all samples (23). The total volumes of lysis and DNA binding buffers were modified to account for the small sample size: 1.7 mL lysis buffer (1.6 mL 0.5 M EDTA (0.5M); 100 µL SDS (10 %); and 20 µL proteinase K (20mg/ml)) and 3 mL guanidinium DNA binding buffer. Negative extraction blank controls were included at a ratio of 1:7, control per calculus samples. Next, shotgun libraries were generated without enzymatic damage repair (23). Briefly, 20 µL of DNA extract was prepared by enzymatic polishing to produce blunt ended fragments prior to ligation of truncated barcoded Illumina adaptors. To maintain sequence complexity, each sample was amplified in triplicate using 13 cycles with HiFi Taq polymerase and full length indexed Illumina adaptors (26). The resulting triplicate amplifications were pooled and purified with the Agencourt AMPure XP system. All samples were then pooled, purified, and then quantified using a TapeStation (Agilent) and quantitative PCR (KAPA Illumina quantification kit) to create a 2 nM sequencing library. In total, 128 of the 161 (79.5%) dental calculus samples yielded high-quality DNA libraries that were sequenced using a high output 2 × 150 bp kit on the Illumina NextSeq.

### Bioinformatic analysis and quality filtering

To identify the microbial communities preserved within samples, raw FastQ files were demultiplexed using sabre (https://github.com/najoshi/sabre). Bbmerge (http://sourceforge.net/projects/bbmap/) was used to merge reads (5 bp overlap), and AdapterRemoval (27) was applied to identify and remove the 5’ and 3’ barcode and adaptor sequences. Reads greater than 300 bp were discarded, as they likely represent modern contamination (15). Microbial species were identified using MALTX (23,28) against the NCBI nr database (2014), and the resulting information was uploaded and filtered using default LCA parameters in MEGAN5 (29). The identified reads within the samples were then normalized to 129,760 sequences, which was the lowest number of reads observed in any sample. Lastly, laboratory contaminant signal was removed from all samples by filtering any contaminant species observed in the negative controls from the calculus samples, as previously described (23). 464 contaminant taxa (26.4% of all species level identifications) were removed from the overall dataset.

### Statistical analysis of associated metadata

The oral geography of the samples and the workflow metadata were compared to the microbiota to assess the potential impact of oral geography and sample handling. All sequence reads that could be assigned to organisms for each sample were exported from MEGAN5 and transformed for use in QIIME (V1.8) (30). PERMANOVA tests (anosim; 9,999 permutations) were used to correlate oral sampling location (*e.g.* upper or lower jaw, tooth type, and tooth surface), sample information (*i.e.* fragment size and sub- or supra-gingival), and processing details (*i.e.* date of sampling, extraction, sequencing) with the Bray-Curtis matrix (Table S2). Following this analysis, samples were filtered based on tooth type, tooth surface, and sub- or supragingival calculus. The three largest datasets were taken forward for further analysis: Molar, Lingual/Palatal, Supragingival (n = 36); Premolar, Lingual/Palatal, Supragingival (n = 18); and Incisor, Lingual/Palatal, Supragingival (n = 18). These groupings were then utilized to examine metadata associated with health, socio-cultural, and environmental factors that drive microbiota diversity using a PERMANOVA test (anosim) in QIIME with a cutoff of p = 0.05 for significance. A G-test test was used to identify specific species that contributed to the differences observed in each metadata category with a cutoff of p = 0.05 for significance. To ensure statistical tests were appropriate, a minimum of five samples per group was used when comparing two metadata categories, while a minimum of three samples per group was enforced when three or more metadata categories were compared. Samples without metadata for a certain category were excluded from those specific tests.

## Results

### Robust oral microbiota recovered from historical London samples

Across the nine archaeological sites included in this study, all samples were dominated by bacterial phyla typical of the modern and ancient human mouth, including Actinobacteria, Firmicutes, and Bacteroidetes (22,23) (Figure 2), consistent with recovery of an oral community structure in these samples. We then examined the impacts of contaminant DNA and laboratory methods on these data. To identify ancient calculus samples that lack true endogenous signal, we looked to identify a signal shared among negative controls that is representative of a laboratory environment. Bray-Curtis dissimilarities were calculated in MEGAN5 for the published ancient calculus control (n=1) (23) and negative controls sequenced within this study (n=10). Negative controls with Bray-Curtis dissimilarities for all negative controls were <0.6 from each other, indicating a consistent microbial signal present in the laboratory. Any calculus sample that fell within four times the mean standard deviation of this group was considered poorly preserved (*i.e.* largely consisted of laboratory contamination) and not fit for downstream analysis. Lastly, PERMANOVA tests indicated that there was no correlation between the microbial community structure of all samples and the date of sampling, DNA extraction, or library preparation (anosim; p > 0.01; Table S2). Overall, 128 dental calculus samples from Medieval London appear to contain robust oral microbiota signatures and so were retained for downstream analysis.

**Figure 2:**
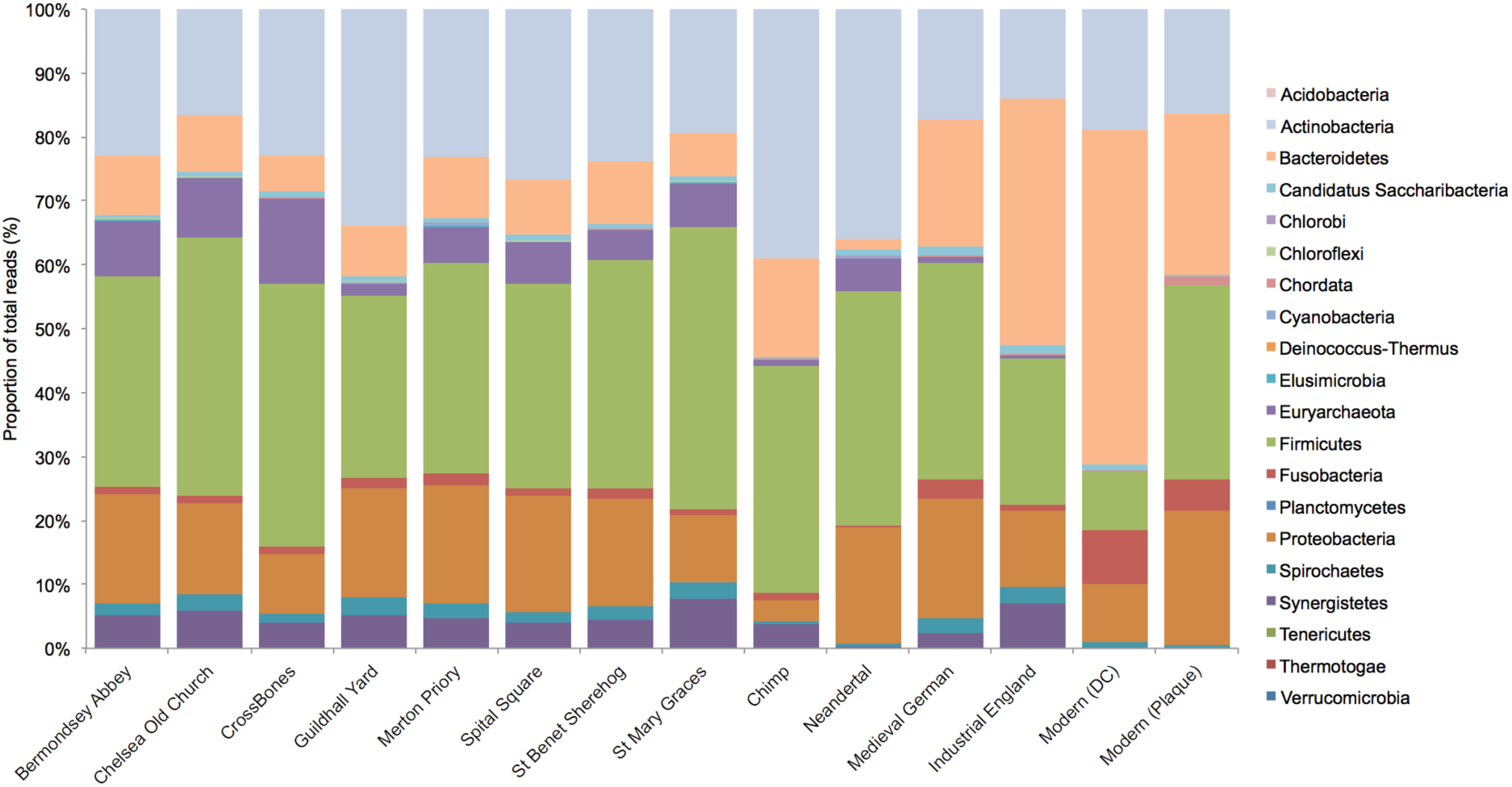
Stacked bar chart showing Museum of London samples (grouped by archaeological site) and previously analysed samples from Weyrich *et al.* (2017) and Warinner *et al.* (2014) and modern plaque from Belda-Ferre, P. et al (2012)

### Oral geography accounts for some variation between individuals

To examine if oral geography impacts the microbiota present in ancient dental calculus samples, we correlated oral microbial diversity in all ancient samples (n = 128) with the oral sampling location. The tooth type was significantly correlated with the microbiota in each sample (anosim; p=0.0001; Table S2). Tooth type was also the variable that explained the most variation on the PCoA plot calculated from Bray-Curtis values of all samples (Figure 3), accounting for 44.9 % of the variation in the data. We then controlled for tooth type by subsequently processing the samples from each tooth type independently. Within tooth types, we observed that tooth surface (buccal, lingual/palatal, interproximal) was a significant driver of diversity (anosim; p < 0.01; Figure 4; Table S2), except for canines. In the canine data set, samples correlated with lab extraction date and library ID (anosim; p = 0.0286 and p = 0.0105, respectively), indicating potential introduction of background laboratory contaminants during sample preparation. However, the canine data set also contained the fewest samples (n=14), so this correlation may be associated with insufficient sampling depth. For stringency, canine samples were not included in further analysis. This highlights the need to record laboratory information for data processing. Lastly, we examined differences between sub- and supragingival microbiota, as differences between these sampling locations have been reported in modern populations (31). Unfortunately, there were very few samples of subgingival dental calculus in our study (5 of 128 samples; Table S1) limiting our ability to test this. However, to minimize any contributions from this potential bias, the five subgingival samples were removed from downstream analysis. Overall, these data indicate that oral geography can contribute to microbiota diversity within ancient calculus and needs to be controlled for in order to avoid introducing bias.

**Figure 3:**
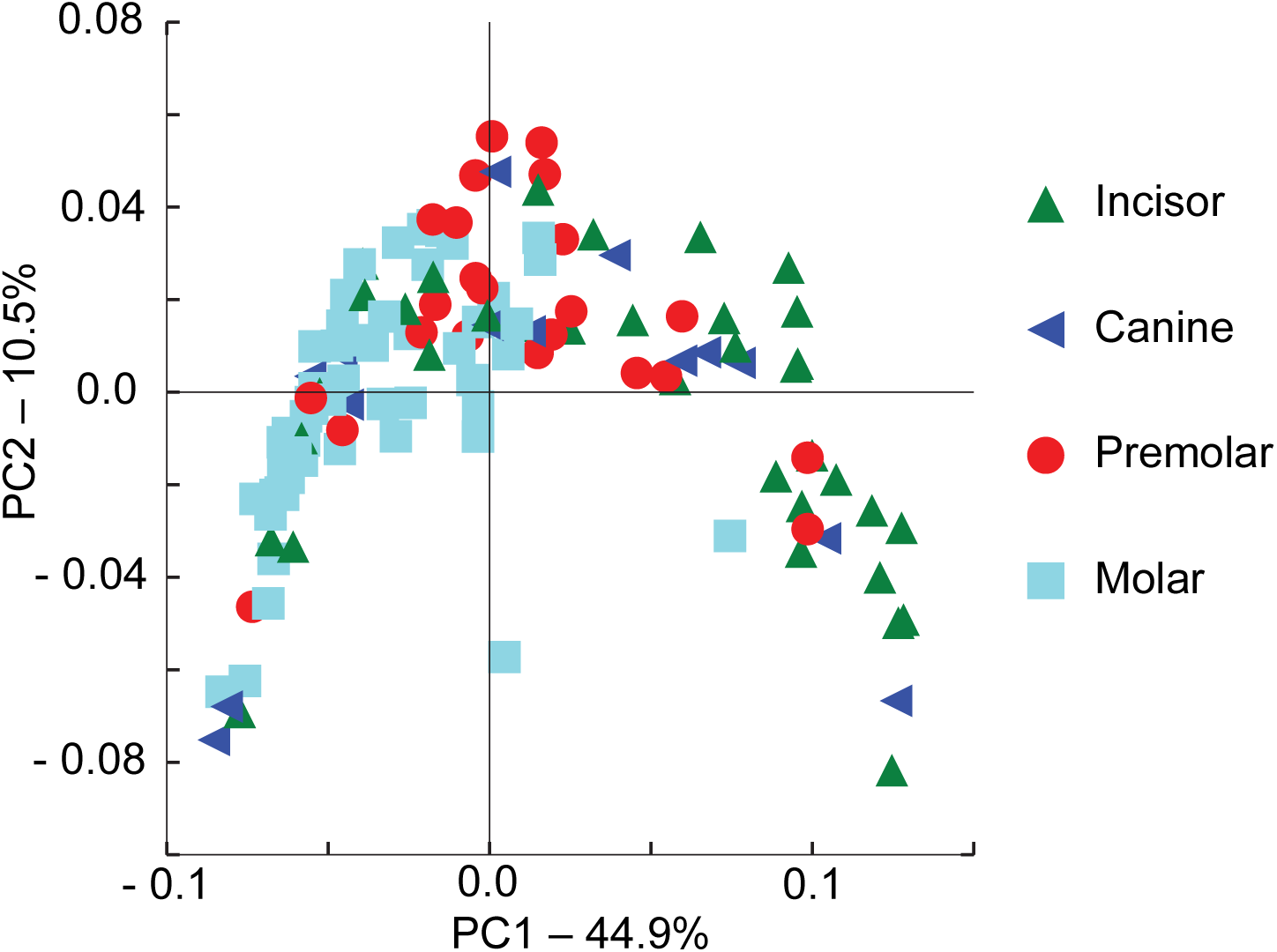
PCoA plot of Bray-Curtis dissimilarity for all dental calculus samples (n = 128). Individual samples are labeled by tooth type. The first axis (PC1) indicates the separation of tooth types, notably molar and incisor.

**Figure 4:**
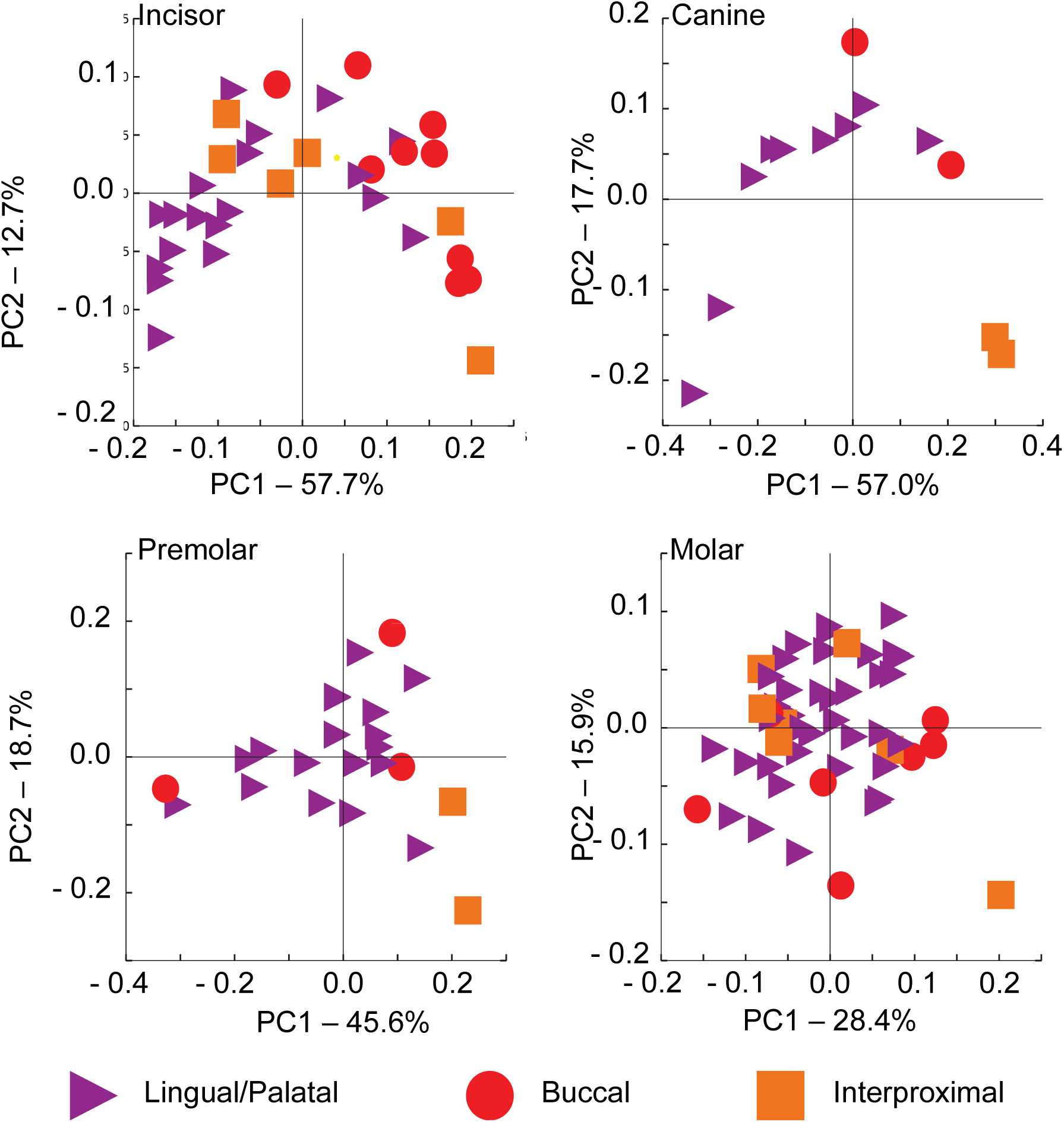
PCoA plot of Bray-Curtis dissimilarity for all dental calculus samples divided by tooth type. Individual samples are coded by the tooth surface from which the sample was taken.

### Microbiota are correlated with disease

We examined links between ancient oral microbiota and evidence of oral and systemic diseases, a wide-range of lifestyle factors, and temporal differences. First, we examined relationships between microbiota (supragingival dental calculus samples within three data sets: the lingual/palatal side of the molar (n=36), premolar (n=18), and incisor (n=18) teeth) and sample metadata, including archaeological site, paleopathology, cultural factors, disease, and period (Table S1). There was no association between microbiota and archaeological site or time period (anosim; p= <0.01; Tables S4-6), which have been assumed to be driving factors in previous aDNA studies (18,23). In addition, no metadata category significantly explained variation within the incisor group (anosim; p= <0.01; Table S4). We also examined factors previously hypothesized to drive microbiota differences in ancient populations, including site location (18) and diet (14), and those modern studies indicate may be present in ancient populations, *e.g.* rural-urban status (32); none were found to be significant. However, disease factors were related to variation within the molar and premolar groups. Abscesses (oral disease; p = 0.0052) and various bone pathologies (*i.e.* porosity, osteophytic lipping, and non-specific periostitis – examples of systemic disease; p = 0.0308 – 0.0466) were all significantly linked with microbiota variation within the molar data set (Table S6), and these correlations were investigated further.

### Oral disease associations with microbiota alteration

The presence of abscesses within the oral cavity was correlated with microbiota within supragingival lingual/palatal dental calculus of molar teeth (p < 0.0052; Table S6). Therefore, we explored the microbial species linked to this oral disease. 30 bacterial species were increased in individuals with dental abscesses, while 20 species were decreased (G-test, p < 0.05; Table S7). Notably, *Prevotella* and *Streptococcus* species, which are associated with dental abscesses in modern humans (33), were significantly increased in the ancient individuals with abscesses (178.0% and 330.0% of the mean read count in healthy individuals, respectively; Table S7). However, other species linked with abscess in modern populations, such as *Porphymonas gingivalis* and *Treponema denticola* (34), were found to be significantly lower in ancient individuals with abscesses (51.0% and 62.2%, respectively). In addition, other potential microbes involved in abscess formation were not significantly different (*e.g. Clostridium*, *Fusobacterium*, and *Bacteroides* species). Individuals with dental abscesses also had significantly increased proportions of archaea, including *Methanobrevibacter* species, which are primary colonizers of the tooth surface (23) and are associated with severe periodontal disease in modern populations (35) (185.5%; Table S7). Together, this suggests that dental abscesses present in ancient Londoners were polymicrobial, similar to modern populations, and left unique signatures in ancient oral microbiota, including the presence of archaea.

### Relationships between bone pathologies and microbiota composition

#### Porosity and Osteophytic lipping

Observations of bone porosity indicate excessive opening of the pores in bone to allow blood, nerve, and other soft tissue to enter (36). This can be associated with age of the individual (37), as well as specific diseases such as anemia and rickets (38). Osteophytic lipping (outgrowths of bone) form around the joint surfaces and are often present alongside arthritis (39). Within our molar data set, all individuals with bone porosity (n = 9) also had osteophytic lipping in joints, while five additional individuals only had osteophytic lipping (Table S1). Consequently, taxonomic differences may not be specifically associated with either trait. Porosity was linked to an increase in 28 microbiota taxa and a decrease in 28 taxa (G-test; p < 0.05; Table S8). *Prevotella* and *Methanobrevibacter* species, which have previously been associated with bone diseases, such as arthritis (40), were increased in individuals with bone porosity. However, as with abscesses, species linked to oral inflammatory diseases and generally increased systemic inflammation, such as *Porphymonas gingivalis, Treponema denticola,* and *Tannerella forsythia* (6,34), were decreased in individuals with bone porosity (24.6%, 72.3%, and 65.6% respectively). Osteophytic lipping was associated with an increase in 19 microbiota taxa (G-test; p < 0.05, Table S9). These taxa are similar to those increased with porosity; for example, *Prevotella* species and *Methanobrevibacter* species are also present in higher abundance in individuals with osteophytic lipping (145.7% and 290.1%, respectively). Together, these data suggests that specific oral microbiota communities are found in ancient Londoners with joint disease.

#### Periostitis

Periostitis is an inflammation of tissue surrounding the bone and often indicative of disease in the underlying bone (41). In living patients, it can be associated with various diseases (including syphilis and skin ulcers) and is linked with trauma (41). Within calculus samples from individuals with periostitis, *Prevotella* species and *Methanobrevibacter* species were again increased (147.8% and 217.7% respectively; Table S10), as observed with porosity and osteophytic lipping. However, several unique bacterial species were linked with this disease, including *Capnocytophaga*, *Clostridia*, *Eubacterium*, *Firmicutes*, *Mogibacterium*, *Neisseria*, and *Pyramidobacter* (G-test; p < 0.05; Table S10). This indicates that periostitis may be linked to a unique shift in the human microbiota, which is different to those observed for porosity and osteophytic lipping.

#### Schmorl’s nodes

In contrast to the molar samples, the microbiota of premolar teeth microbiota were correlated with Schmorl’s nodes – a vertebral bone pathology (anosim; p = 0.05; Table S5). While Schmorl’s nodes are common, their pathological status is poorly understood. It is thought that they appear in response to herniation of the cartilaginous disc into the vertebral body as a result of direct physical stress (42,43) Methanobrevibacter was increased in individuals with the pathology (129.3%; Table S11). This is consistent with the previous pathologies examined. In contrast, Prevotella was decreased in those with Schmorl’s Nodes (75.1%; Table S11). It is worth noting that while each of these bone pathologies is classified independently, it is possible that they have arisen from a similar cause, such as an increase in manual labor within this population (44). The lack of a specific causal factor for these paleopathology could explain why similar associations with the oral microbiota are inferred.

## Discussion

In the largest ancient dental calculus study to date, we investigated the roles that experimental biases can play in ancient dental calculus research and revealed direct associations between ancient human oral and systemic health and the oral microbiota. We show that the oral geography of the mouth is a critical factor when identifying fine-scale factors that impact the human microbiota, as tooth type and tooth surface explained the greatest amount of variation observed in our data set. Once these and other technical factors are controlled, information that underpins the evolutionary history of the oral microbiota can be examined, highlighting the potential of dental calculus to study the impacts of cultural and environmental factors on the oral microbiota and, thus, the evolutionary history of the modern oral microbiota.

Within this study, the oral geography of dental calculus accounted for the most significant source of variation within the data, raising questions about conclusions drawn from previous ancient calculus research. Several studies have combined dental calculus samples from different regions within the mouth (22) or not factored in tooth type or tooth surface (23). Our study suggests that there are limitations to these approaches and that ancient dental calculus studies moving forward should account for tooth type and surface when examining correlations between individual metadata and oral microbiota. Even though it markedly reduced the total sample size per group, intrapersonal variation largely masked many of the links between disease and microbiota. Notably, different sampling locations within the mouth also revealed individual associations with skeletal metadata. While limited sample numbers in some tooth groupings may affect these findings, oral microbial communities from different teeth appear to respond differently in states of disease. In future, a study of the oral geography within single ancient individuals, improved bioinformatic methodologies, and increased sample sizes will help understand and account for intrapersonal variation.

This study also highlights additional technical factors that may influence ancient oral microbiota diversity, including contamination and experimental metadata. To address potential laboratory contamination, we compared samples to negative blank controls to confirm that samples did not have a microbial signal consistent with the laboratory environment. Conservatively, any microbial species in common with the negative controls was also removed from the samples. Samples were also randomized before undergoing laboratory protocols. This demonstrates the care with which ancient microbiota data must be handled, as researchers must be careful to collect, document, and analyse sampling and experimental metadata to accurately interpret findings in ancient microbiota. In future studies, the collection of detailed metadata will improve the identification of local cultural and environmental factors that influenced the microbiota, and scientists will be better able to identify potential links between oral microbiota and a past individual’s life history.

This is the first study to examine ancient oral microbiota variation with respect to a wide-range of individualistic metadata, including sex, age, rural-urban status, religion, time, and location. Despite this wealth of information, this study revealed that disease appears to be the single largest determinant of oral microbiota variation in ancient London. We identified alterations in community structure associated with both oral and systemic disease. In modern populations, oral microbiota have been linked to a wide range of oral and systemic diseases, as diverse as dental caries, gum disease, arthritis, heart disease, pre-term birth, Alzheimer’s disease, and mental disorders (5,6,9,34,45). Although past aDNA studies have identified ancient oral pathogens as potential signatures of disease (18,21,22), we identify community level changes linked to abscesses that include both known, abscess-associated pathogens (*Prevotella* and *Streptococcus* species) and lesser known species (*Methanobrevibacter* species). In addition, microbiota from molar teeth correlated with bone disease, such as porosity, osteophytic lipping and peristotitis and, in all cases, were demarcated by an increase in oral *Prevotella* species. In modern populations, *Prevotella* species are associated with arthritis (6), although the mechanism underlying this link is not yet known. There may be a direct link between this species and Porosity, Oesteophytic Lipping, and Periostitis because oral bacteria can escape the mouth and cause inflammation and lesions elsewhere in the body (46). Alternatively, *Prevotella* species or community structure may serve as an oral marker for overall community alterations that are linked to generalized inflammation in the body. Consequently, causation cannot be inferred, and further studies are needed to investigate mechanistic relationships.

Bone diseases linked to greater physical stress and malnutrition are typically seen in ancient individuals with low socio-economic rank (47). In London, low socioeconomic status has been previously associated with poor bone health (47). While it is possible that these bone diseases directly led to alterations in microbiota, it is also possible that these relationships between disease and microbiota are explained by greater lifestyle factors, such as low socioeconomic status. In fact, several studies to date have linked socioeconomic status to microbiota community composition (48–50). Our results are consistent with these observations, and may suggest that oral microbiota can be used as a marker for socioeconomic status in the past. Further research is needed to investigate this trend in different human cultures worldwide.

Large-scale sampling of ancient dental calculus from a single population has provided an unprecedented opportunity to reveal information about how past lifestyles shape the future of human health. This study reveals the impacts of technical factors and highlights how critical disease was in the past for shaping the evolutionary history of human oral microbiota. In addition, this study provides fundamental baseline evidence to suggest that the oral microbiota can be used as archaeological biomarkers of cultural, health, and environmental change, allowing researchers to gain new and more detailed insights when other archaeological information is unavailable.

## Author’s Contributions

AGF, JB, RR, AC, KD and LSW contributed to study design. AGF, JB, and RR submitted destructive sampling applications, collated metadata, and collected samples. AGF conducted laboratory work and bioinformatic analysis. AGF, NG, KD, AC, and LSW contributed to data interpretation. AGF and LSW outlined and wrote the manuscript, and KD, AC, and LSW edited the manuscript.

## Acknowledgements

We would like to thank the Museum of London for allowing us to collect and destructively analyse ancient dental calculus samples from the specimens utilized in this study.

## References

1. Rosenbaum M, Knight R, Leibel RL. The gut microbiota in human energy homeostasis and obesity. Trends Endocrinol Metab. 2015 Sep;26(9):493–501.

2. Trivedi B. Microbiome: The surface brigade. Nature. 2012;492(7429):2.

3. Li X, Wang J, Joiner A, Chang J. The remineralisation of enamel: a review of the literature. J Dent. 2014 Jun;42, Supplement 1:S12–20.

4. Iwase T, Uehara Y, Shinji H, Tajima A, Seo H, Takada K, et al Staphylococcus epidermidis Esp inhibits Staphylococcus aureus biofilm formation and nasal colonization. Nature. 2010 May 20;465(7296):346–9.

5. Noverr MC, Huffnagle GB. Does the microbiota regulate immune responses outside the gut? Trends Microbiol. 2004 Dec; 12(12):562–8.

6. Rautemaa R, Lauhio A, Cullinan MP, Seymour GJ. Oral infections and systemic disease—an emerging problem in medicine. Clin Microbiol Infect. 2007 Nov 1;13(11):1041–7.

7. Weyrich LS, Feaga HA, Park J, Muse SJ, Safi CY, Rolin OY, et al Resident Microbiota Affect Bordetella pertussis Infectious Dose and Host Specificity. J Infect Dis [Internet]. 2013 Dec 15; Available from: http://www.ncbi.nlm.nih.gov/pubmed/24227794

8. Farrell JJ, Zhang L, Zhou H, Chia D, Elashoff D, Akin D, et al Variations of oral microbiota are associated with pancreatic diseases including pancreatic cancer. Gut. 2012 Apr 1;61(4):582–8.

9. Grau AJ, Becher H, Ziegler CM, Lichy C, Buggle F, Kaiser C, et al Periodontal Disease as a Risk Factor for Ischemic Stroke. Stroke. 2004 Feb 1;35(2):496–501.

10. Turnbaugh PJ, Ley RE, Mahowald MA, Magrini V, Mardis ER, Gordon JI. An obesity-associated gut microbiome with increased capacity for energy harvest. Nature. 2006 Dec 21;444(7122):1027–31.

11. David LA, Maurice CF, Carmody RN, Gootenberg DB, Button JE, Wolfe BE, et al Diet rapidly and reproducibly alters the human gut microbiome. Nature. 2014 Jan 23;505(7484):559–63.

12. Chen YE, Tsao H. The skin microbiome: Current perspectives and future challenges. J Am Acad Dermatol. 2013 Jul;69(1):143-155.e3.

13. Lax S, Smith DP, Hampton-Marcell J, Owens SM, Handley KM, Scott NM, et al Longitudinal analysis of microbial interaction between humans and the indoor environment. Science. 2014 Aug 29;345(6200):1048–52.

14. Weyrich LS, Dobney K, Cooper A. Ancient DNA analysis of dental calculus. J Hum Evol. 2015 Feb;79:119–24.

15. Weyrich LS. Evolution of the Human Microbiome and Impacts on Human Health, Infectious Disease, and Hominid Evolution. In: Gontier N, editor. Reticulate Evolution [Internet]. Springer International Publishing; 2015 [cited 2015 Oct 27]. p. 231–53. (Interdisciplinary Evolution Research). Available from: http://link.springer.com/chapter/10.1007/978-3-319-16345-1_9

16. Warinner C, Speller C, Collins MJ. A new era in palaeomicrobiology: prospects for ancient dental calculus as a long-term record of the human oral microbiome. Phil Trans R Soc B. 2015 Jan 19;370(1660):20130376.

17. Warinner C, Speller C, Collins MJ, Lewis Jr. CM. Ancient human microbiomes. J Hum Evol. 2015 Feb;79:125–36.

18. Adler CJ, Dobney K, Weyrich LS, Kaidonis J, Walker AW, Haak W, et al Sequencing ancient calcified dental plaque shows changes in oral microbiota with dietary shifts of the Neolithic and Industrial revolutions. Nat Genet. 2013 Apr;45(4):450–5.

19. White DJ. Processes contributing to the formation of dental calculus. Biofouling. 1991 Aug 1;4(1–3):209–18.

20. Dobney K, Brothwell D. Dental Calculus: Its Relevance to Ancient Diet and Oral Ecology. In: Cruwys E, Foley RA, editors. Teeth and Anthropology. 1986. p. 55–81. (British Archaeological Reports).

21. De La Fuente C, Flores S, Moraga M. DNA from Human Ancient Bacteria: A Novel Source of Genetic Evidence from Archaeological Dental Calculus. Archaeometry. 2013;55(4):767–78.

22. Warinner C, Rodrigues JFM, Vyas R, Trachsel C, Shved N, Grossmann J, et al Pathogens and host immunity in the ancient human oral cavity. Nat Genet. 2014;

23. Weyrich LS, Duchane S, Soubrier J, Arriola L, Llamas B, Breen J, et al Reconstructing Neandertal behavior, diet, and disease using ancient DNA from dental calculus. Nature. 2016;

24. Simón-Soro Á, Tomás I, Cabrera-Rubio R, Catalan MD, Nyvad B, Mira A. Microbial Geography of the Oral Cavity. J Dent Res. 2013 Jul 1;92(7):616–21.

25. Salter SJ, Cox MJ, Turek EM, Calus ST, Cookson WO, Moffatt MF, et al Reagent and laboratory contamination can critically impact sequence-based microbiome analyses. BMC Biol. 2014 Nov 12;12(1):87.

26. Meyer M, Kircher M. Illumina Sequencing Library Preparation for Highly Multiplexed Target Capture and Sequencing. Cold Spring Harb Protoc. 2010 Jun 1;2010(6):pdb.prot5448.

27. Lindgreen S. AdapterRemoval: easy cleaning of next-generation sequencing reads. BMC Res Notes. 2012;5:337.

28. Herbig A, Maixner F, Bos KI, Zink A, Krause J, Huson DH. MALT: Fast alignment and analysis of metagenomic DNA sequence data applied to the Tyrolean Iceman. bioRxiv. 2016 Apr 27;050559.

29. Huson DH, Auch AF, Qi J, Schuster SC. MEGAN analysis of metagenomic data. Genome Res. 2007 Mar 1;17(3):377–86.

30. Caporaso JG, Kuczynski J, Stombaugh J, Bittinger K, Bushman FD, Costello EK, et al QIIME allows analysis of high-throughput community sequencing data. Nat Methods. 2010 May;7(5):335–6.

31. Zijnge V, Leeuwen MBM, van Degener JE, Abbas F, Thurnheer T, Gmür R, et al Oral Biofilm Architecture on Natural Teeth. PLOS ONE. 2010 Feb 24;5(2):e9321.

32. Ayeni FA, Biagi E, Rampelli S, Fiori J, Soverini M, Audu HJ, et al Infant and Adult Gut Microbiome and Metabolome in Rural Bassa and Urban Settlers from Nigeria. Cell Rep. 2018 Jun 5;23(10):3056–67.

33. Shweta, Prakash SK. Dental abscess: A microbiological review. Dent Res J. 2013;10(5):585–91.

34. Rocas IN, Siqueira JF Jr, Santos KR, Coelho AM. “Red complex” (Bacteroides forsythus, Porphyromonas gingivalis, and Treponema denticola) in endodontic infections: a molecular approach. Oral Surg Oral Med Oral Pathol Oral Radiol Endod. 2001 Apr;91(4):468–71.

35. Horz H-P, Conrads G. Methanogenic Archaea and oral infections — ways to unravel the black box. J Oral Microbiol [Internet]. 2011 Feb 23 [cited 2016 Feb 17];3. Available from: http://www.ncbi.nlm.nih.gov/pmc/articles/PMC3086593/

36. Cardoso L, Fritton SP, Gailani G, Benalla M, Cowin SC. Advances in assessment of bone porosity, permeability and interstitial fluid flow. J Biomech. 2013 Jan 18;46(2):253–65.

37. Gabet Y, Bab I. Microarchitectural Changes in the Aging Skeleton. Curr Osteoporos Rep. 2011 Sep 8;9(4):177.

38. Pinhasi R, Mays S. Advances in Human Palaeopathology. John Wiley & Sons; 2008. 410 p.

39. Ak J. Osteophytes and the osteoarthritic femoral head. J Bone Joint Surg Br. 1975 1975;57(3):314–24.

40. Scher JU, Abramson SB. The microbiome and rheumatoid arthritis. Nat Rev Rheumatol. 2011 Aug 23;7(10):569–78.

41. Ortner DJ. Identification of Pathological Conditions in Human Skeletal Remains. Academic Press; 2003. 663 p.

42. Kyere KA, Than KD, Wang AC, Rahman SU, Valdivia–Valdivia JM, Marca FL, et al Schmorl’s nodes. Eur Spine J. 2012 Apr 28;21(11):2115–21.

43. Sonne-Holm S, Jacobsen S, Rovsing H, Monrad H. The epidemiology of Schmorl’s nodes and their correlation to radiographic degeneration in 4,151 subjects. Eur Spine J. 2013 Mar 16;22(8):1907–12.

44. Black J. London: a history. Carnegie; 2009.

45. Selwitz RH, Ismail AI, Pitts NB. Dental caries. The Lancet. 2007 Jan 12;369(9555):51–9.

46. Billing F. Focal Infection: Its broader application in the etiology of general disease. J Am Med Assoc. 1914;63(11):5.

47. Newman SL, Gowland RL. Dedicated Followers of Fashion? Bioarchaeological Perspectives on Socio-Economic Status, Inequality, and Health in Urban Children from the Industrial Revolution (18th–19th C), England. Int J Osteoarchaeol. 2017 Mar 1;27(2):217–29.

48. Kau AL, Ahern PP, Griffin NW, Goodman AL, Gordon JI. Human nutrition, the gut microbiome and the immune system. Nature. 2011 Jun;474(7351):327–36.

49. Belstrøm D, Holmstrup P, Nielsen CH, Kirkby N, Twetman S, Heitmann BL, et al Bacterial profiles of saliva in relation to diet, lifestyle factors, and socioeconomic status. J Oral Microbiol. 2014 Jan 1;6(1):23609.

50. Miller GE, Engen PA, Gillevet PM, Shaikh M, Sikaroodi M, Forsyth CB, et al Lower Neighborhood Socioeconomic Status Associated with Reduced Diversity of the Colonic Microbiota in Healthy Adults. PLOS ONE. 2016 Feb 9;11(2):e0148952.

